# HADDOCK3: A modular and versatile platform for integrative modelling of biomolecular complexes

**DOI:** 10.1101/2025.04.30.651432

**Authors:** Marco Giulini, Victor Reys, João M. C. Teixeira, Brian Jiménez-García, Rodrigo V. Honorato, Anna Kravchenko, Xiaotong Xu, Raphaëlle Versini, Anna Engel, Stefan Verhoeven, Alexandre M.J.J. Bonvin

## Abstract

HADDOCK is a widely used resource for integrative modelling of a variety of biomolecular complexes that is able to incorporate experimental knowledge into physics-based calculations during complex prediction, refinement, scoring and analysis.

Here we introduce HADDOCK3, the new modular version of the program, in which the original, parameterisable albeit rigid pipeline has been first broken down in a catalogue of independent modules and then enriched with powerful analysis tools and third-party integrations. Thanks to this increased flexibility, HADDOCK3 can now handle multiple integrative modelling scenarios, providing a valuable, physics-based tool to enrich and complement the predictions made by machine learning algorithms in the post-AlphaFold era. We present examples of successful applications of HADDOCK3 that were not feasible with the previous versions of HADDOCK, highlighting its expanded capabilities.

The HADDOCK3 software source code is freely available from the GitHub repository (https://github.com/haddocking/haddock3) and comes with an online user guide (www.bonvinlab.org/haddock3-user-manual). All example data described in this manuscript are available at https://github.com/haddocking/haddock3-paper-data.

## Introduction

Modelling biomolecular interactions plays a central role in structural biology, as most biological processes take place through intricate networks of interactions occurring in the crowded cellular and extracellular environments.

In the last ten years, much progress has been made in the computational modelling of such interactions, in particular thanks to the development of deep learning-based methods able to accurately predict the three-dimensional structure of protein-protein complexes from sequences [1, 2, 3, 4, 5], recently expanded to the modelling of nucleic acids [6], as well as lipids, small molecules, and post-translational modifications [7, 8].

Classical approaches such as molecular docking are complementary to machine learning architectures [9, 10], as they use physics-based methods to tackle challenging systems that cannot be accurately modelled using current machine learning methods. Although the typical success rate of these approaches is quite low when used in *ab-initio* mode (i.e. with no apriori information), it can be substantially improved in data-driven mode, i.e., when some experimental knowledge is available about the interaction [11, 12, 13].

Incorporating existing experimental information in the modelling process is at the core of the HADDOCK software [11], which uses the concept of ambiguous interaction restraints (AIRs) to bias physics-based docking towards models that maximize the consistency with the input data.

HADDOCK has been widely used in the last twenty years, with more than 60 000 registered users being able to perform almost 700 000 docking runs on the HADDOCK webserver [9] which has been in operation since 2008. The HADDOCK2.X series of software [14, 15, 9] consists of a rigid, albeit highly parameterisable pipeline, in which rigid-body docking is followed by semi-flexible interface refinement and a final energy minimisation. Optionally, the final minimisation can be replaced by a refinement in explicit solvent.

Here we introduce HADDOCK3, the next version of the HADDOCK software, which represents a complete rewriting and rethinking of the entire architecture. The original HADDOCK pipeline has been broken down into a catalogue of independent modules that can be, in large parts, freely combined into user- and system-specific custom workflows.

This modular and flexible architecture allows HADDOCK3 to handle a multitude of integrative modelling scenarios, providing a valuable, physics-based tool to enrich and complement the predictions made by machine learning algorithms in the post-AlphaFold era. In the following, we describe this new HADDOCK3 version and illustrate its capabilities with a few relevant examples, highlighting how workflows that were not feasible with the previous version can provide accurate structural models.

HADDOCK3 is an open-source software suite, freely available for download from our GitHub repository https://github.com/haddocking/haddock3 or directly from PyPI (https://pypi.org/project/haddock3). The user manual of HADDOCK3 can be accessed at www.bonvinlab.org/haddock3-user-manual.

## Software

### Bringing modularity into HADDOCK

Modelling biomolecular interactions at the structural level typically involves several independent steps: docking the molecules together, refining the obtained models, and scoring them using various scoring functions. HADDOCK3 breaks down the fixed pipeline of the HADDOCK2.X series, thus allowing for a diversification of workflows where not all steps are required and where their order can be customised and optimised for each system under study. This is made possible by HADDOCK3’s modular structure, in which each module is an independent, self-sufficient step to be included in a larger modelling workflow, except for the required initial module that builds the molecular topology of the system. As a result, modules can be freely combined to address different scientific questions.

The modular design also simplifies the integration of new modules into the software, reducing compatibility issues. Name of parameters, their description, and allowed values for each module can be queried using the `haddock3-cfg` command line interface and found online at www.bonvinlab.org/haddock3.

Five categories of modules are currently available in HADDOCK3 (see Table 1):

**Table 1.**
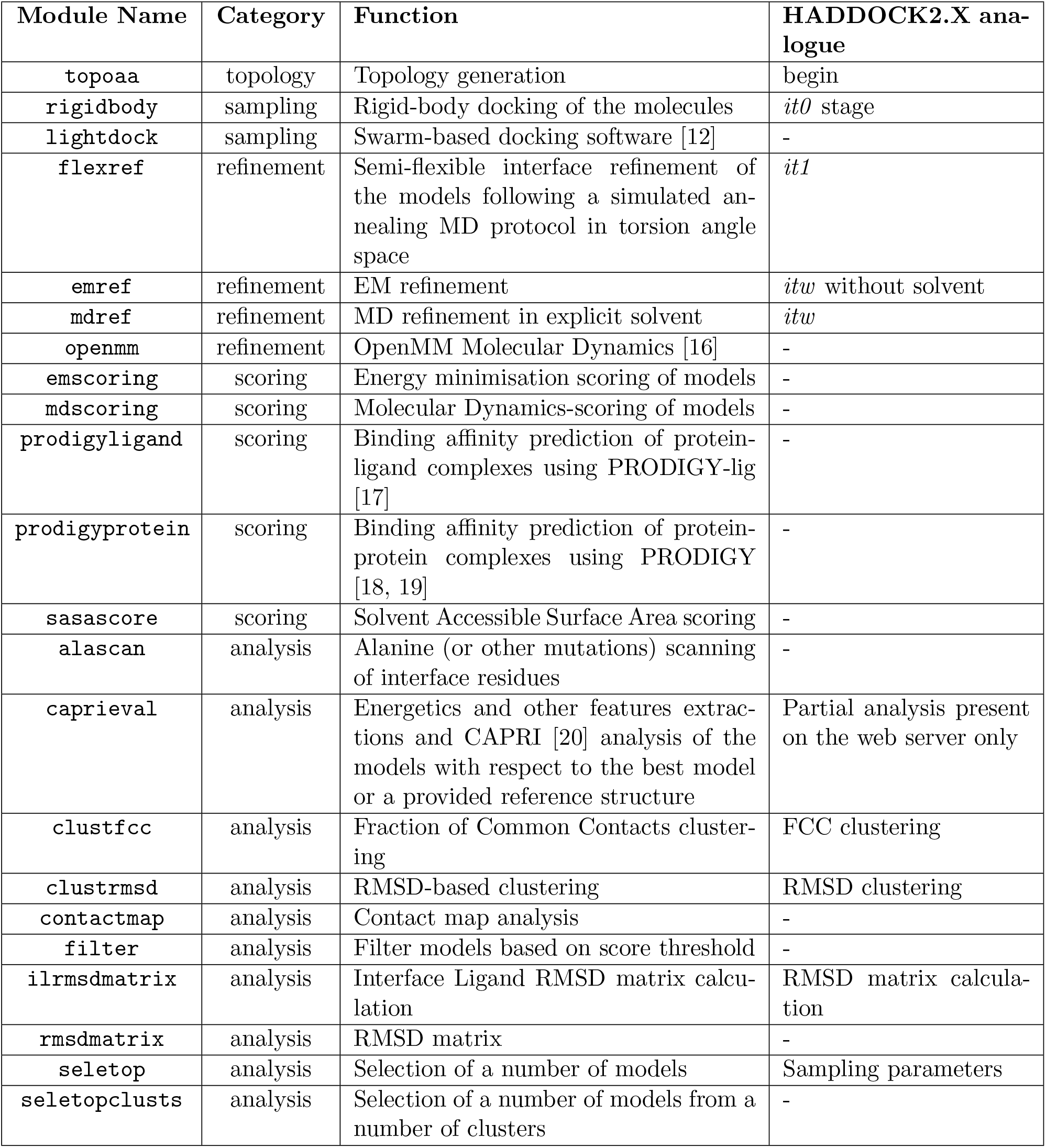
List of the currently available modules in HADDOCK3, each one with its own category, description, and corresponding analogue in the HADDOCK2.x series, when applicable.

- *topology modules*, responsible for the creation of topologies for the available modules;
- *sampling modules*, namely the traditional rigid-body docking methods (including HAD-DOCK2’s *it0* stage, now renamed `rigidbody`);
- *refinement modules*, in which the modelled systems are subject to various refinement algorithms, ranging from simple Energy Minimization (EM) to full explicit solvent Molecular Dynamics (MD) refinement (HADDOCK2’s *it1* now `flexref` and *itw*, now split into `mdref` and `emref`, are part of this category);
- *scoring modules*, in which models are ranked according to various scoring functions;
- *analysis modules*, encompassing a broad class of analysis methods, including model evaluation, clustering and analysis and visualization of interface properties.

### Workflow definition

HADDOCK3 defines workflows using a configuration file compatible with the TOML format but with additional features. In the first part of this file, global parameters are specified, such as the input molecules, the running mode, and the output path. This is followed by the sequence of modules to be executed, each with several parameters that can be fine-tuned or left at their default settings.

Fig. 1 shows the direct translation of the HADDOCK2 pipeline into a HADDOCK3 configuration file. Information to guide the docking is provided to the workflow through ambiguous interaction restraints [11] (`ambig_fname`), ensuring that the resulting models align with the available experimental data. Additionally, unambiguous restraints (`unambig_fname`) can be added to fix the distance between specified pairs of atoms, which is particularly useful when modeling antibody-antigen complexes or dealing with MS crosslink data. The restraint files need to be specified for each module separately, which now gives more flexibility in the definition of which restraints are being used at each stage of the workflow.

**Figure 1:**
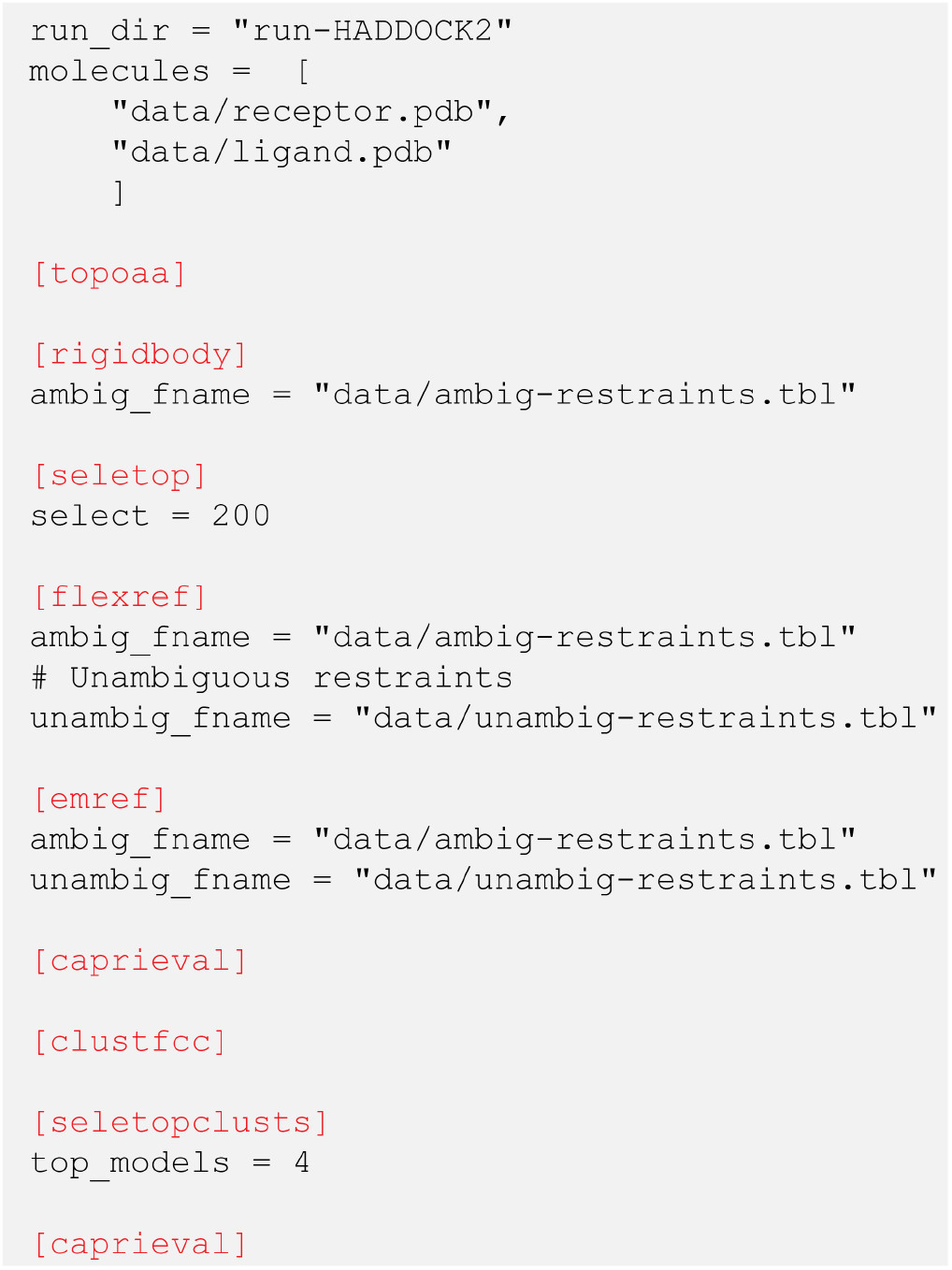
Literal translation of the HADDOCK2 pipeline into an HADDOCK3 workflow. After defining some preliminary global parameters such as the name of the directory where to perform the docking run (`run_dir`) and a list of paths to the input molecules (`molecules`), the list of modules defining the workflow is executed sequentially. First, the allatom topology is built (`topoaa`). Then, the `rigidbody` module (former *it0*) docks the molecules according to the information contained in the ambiguous interaction restraints. A thousand models are generated at this stage by default, but only 200 of them (the ones with the lowest score) are taken to the next step of the workflow (`seletop`), namely the computationally more costly interface refinement (`flexref`). Models are then refined through energy minimization (`emref`) and their statistics extracted through the `caprieval` module, which performs a quality assessment of the models against the top ranked model. Models can be compared instead to a reference structure, if available, by defining the target name. They are then clustered based on the fraction of common contacts [21] (`clustfcc`). `seletopclusts` subsequently selects the top 4 models of each cluster. Lastly, an additional `caprieval` analysis is performed again to obtain cluster-based statistics.

### Workflow execution

HADDOCK3 is a Command Line Interface (CLI) software that requires the user to use a terminal to launch a docking run. Once HADDOCK3 is installed, the path to the workflow configuration file must be passed as input to the `haddock3` CLI to execute a run.

### Workflow results

At execution time, HADDOCK3 will process a workflow in several steps. First, the content of the workflow configuration file is analysed and checked for compliance with the existing modules and parameters. Input file paths are verified, module names and their related parameters are matched against available ones. Once this validation is passed, the run directory is created and input files are stored in the data directory. As a second step, the workflow modules are executed in a stepwise manner. Each module will create its own directory (numbered sequentially) where generated output files will be written.

Upon workflow completion, two post-processing steps are performed by default: first, an analysis phase generates various plots that describe the produced models with their HADDOCK scores and components. Distributions of different metrics used in the Critical Assessment of PRotein Interactions (CAPRI) [20] (ligand-rmsd, interface-ligand-rmsd, fraction of native contacts, and DockQ [22]) are reported, with comparisons made against the best scoring model or a provided reference structure. Then, a traceback step simplifies the analysis of the workflow’s progression, allowing users to trace how models evolved through the various modules and which input conformations or restraint files were used.

### Optimizing execution modes to fit user setups

Being able to run HADDOCK3 on various operating systems and hardware, increasing inter-operability and scale, was a guideline of the development of this new version. HADDOCK3 is written in python3, leveraging an easy installation procedure on most operating systems through `pip install` commands. Detailed installation instructions are provided on the HAD-DOCK3 GitHub repository (https://github.com/haddocking/haddock3). A dedicated parameter (`mode`) enables the user to specify the best execution mode for the system on which the workflow will be run. Three execution modes are available:

- Local mode: HADDOCK3 runs on the current system, using a specified number of cores, which is limited by the available resources. This mode can also be used to, for example, run HADDOCK3 using a full node in a HPC environment.
- Batch mode: HADDOCK3 is initiated from a local server, using the system batch submission system (SLURM or Torque) to submit short jobs to a defined queue.
- MPI mode: This mode allows modelling workflows to span multiple nodes on an HPC architecture, thus enabling distributed computing (provided MPI libraries are installed).

### HADDOCK3 tools

HADDOCK3 comes with several command line interfaces (CLIs), each with its specific goal. The most important ones are described below, but the full list of command line interfaces can be found in the online user manual at www.bonvinlab.org/haddock3-user-manual/clis.html.

- The `haddock3` command is the main CLI allowing to start a HADDOCK3 workflow.
- The `haddock3-cfg` CLI is made to retrieve and list the parameter names, their description, and default values for each available module.
- The `haddock3-analyse` CLI was developed to post-process a docking run and generate tables and plots describing the complexes.
- The `haddock3-restraints` CLI is dedicated at creating and manipulating ambiguous restraints that are used in HADDOCK.
- The `haddock3-score` is a scoring CLI, that seamlessly performs topology generation, short energy minimisation and scoring with the HADDOCK score of the input complex. It bypasses the need of generating a configuration file for the scoring of a single complex.

## Use Cases

In this section we describe four use cases that highlight HADDOCK3 workflows tailored to specific scientific problems that were not possible to address with the previous versions of the software.

### Targeting multiple interfaces

Information-driven modelling of biomolecular complexes requires some knowledge about the interface in order to be effective. In a typical two-body docking scenario, information may be only available for one of the molecules, while the binding site on the partner is ambiguous, meaning that the experimental (or bioinformatics) data suggest several possible binding interfaces.

In previous versions of HADDOCK, two main strategies have been adopted to tackle this quite common scenario. In the first one, two or more separate docking runs are performed, each one targeting a specific interface. A posteriori comparison based on visual inspection and other analysis tools were then used to select the most likely correct binding interface. In the second one, different solvent-accessible patches are defined together as passive (passive in HADDOCK context means that a given residue can be at the interface), hoping that the random removal of restraints and their ambiguity would allow obtaining near-native docking poses. Both strategies are suboptimal, as they rely either on parallel computations and cumbersome additional analysis or on a possibly physically implausible combination of sets of restraints. In HADDOCK3 this problem is addressed by adding the possibility to provide several sets of restraints within a single workflow. The initial rigid body docking models are generated using different sets of restraints. Information about which set was used is propagated for each model to all subsequent stages of the workflow. The scoring function will compare them within the HADDOCK machinery, therefore eliminating the need for multiple parallel runs and a posteriori analysis.

Here we present an example of such an application for the modelling of an antibody-antigen complex, namely the complex (PDB ID 4G6M [24]) between gevokizumab and Interleukin-1*β* (IL-1*β*). Using ARCTIC-3D [23], we automatically extracted from the PDB Knowledge base [25] the five interfaces (epitopes) formed by IL-1*β* with different antibodies, including the one specific to gevokizumab (see Fig. 2a). Using the solvent-exposed residues on the Complementarity-Determining Region (CDR) of the unbound antibody structure (PDB ID 4G6K [24]), we generated five sets of ambiguous interaction restraints, each one effectively targeting a different epitope.

**Figure 2:**
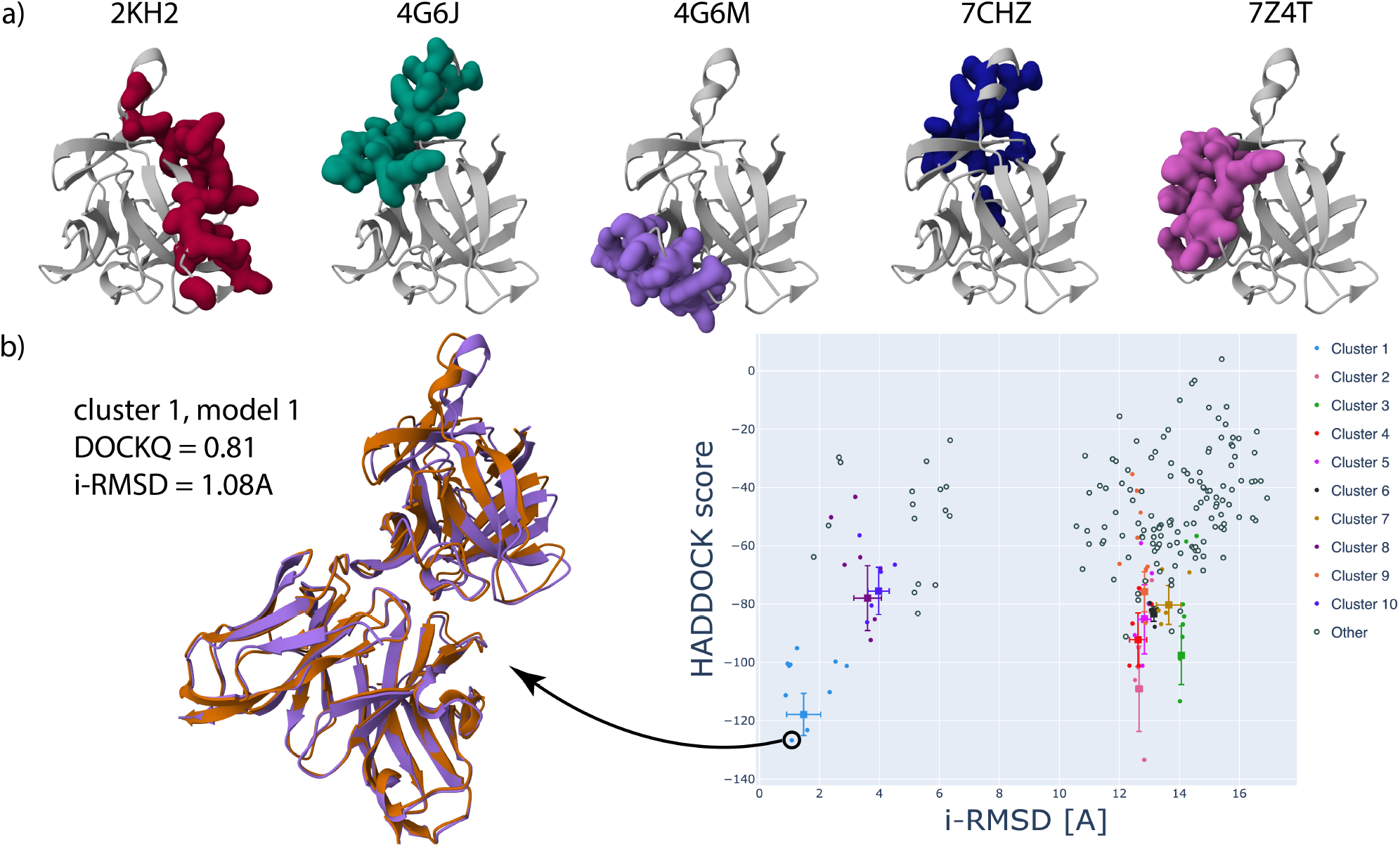
Illustration of multiple restraints targeting different binding sites. a) ARCTIC-3D [23] analysis reveals five different epitopes in Interleukin-1*β*. The PDB ID corresponding to each epitope is reported and the amino acids belonging to it are shown in green on the unbound structure of Interleukin-1*β* (coming from PDB ID 5R8Q, chain B). b) Results of a HADDOCK3 run targeting multiple interfaces within the same workflow. The best model of the top-ranked cluster is shown overlaid on the reference structure (PDB ID 4G6M), highlighting the excellent agreement with the model (DOCKQ = 0.81). On the right, a HADDOCK3 scatter plot between the HADDOCK score and the interface RMSD illustrates how the best cluster can be clearly identified.

A standard HADDOCK workflow is then executed. When considering the cluster-based results, we see how the best cluster is targeting the correct interface and is composed of models of medium-to-high quality following CAPRI criteria, with DockQ values ranging from 0.48 to 0.88 (Fig. 2b). For a comparison, we run a HADDOCK workflow targeting the five different epitopes at the same time (defined together as passive residues, see above). Here HADDOCK cannot generate any acceptable model, with the top-ranked model and overall best model (ranked 136) displaying values of DockQ of 0.034 and 0.072, respectively. In this case, the information encoded in the ambiguous restraints is too sparse and generic, hindering the identification of the correct binding mode of the antibody.

This example demonstrates that HADDOCK3’s ability to handle multiple restraint sets in a single run can effectively identify the correct binding interface with high-quality models.

### Protein-glycan docking: introducing a clustering step after rigid-body docking

Glycans are very flexible biomolecules formed by two or more monosaccharides linked by glycosidic bonds. The chemical composition and branching pattern of such linkages create an incredibly diverse set of possible oligosaccharides [26].

The flexibility provided by HADDOCK3 is particularly suited to model non-covalent protein-glycan interactions, especially when information is available on the protein binding site. In a recent study [27] we demonstrated that tweaking the original HADDOCK recipe provides a substantial increase in the docking performances, reaching a success rate of 60% when considering the top 10 models. Two major ingredients lie at the core of this improvement, namely the increased weight assigned to the hydrophobicity (van der Waals) component of the HADDOCK energy function, and the insertion of a fine-tuned RMSD clustering step between the sampling (`rigidbody` module) step and the flexible interface refinement (`flexref` module). This was not possible in the previous version of HADDOCK due to its pre-defined, fixed pipeline.

We illustrate this slightly more complex workflow in Fig. 3, which we use to model the complex between rainbow trout lysozyme and a linear oligosaccharide (PDB ID 1LMQ [28]). The oligosaccharide consists of three 2-acetamido-2-deoxy-beta-D-glucopyranose and one 2-acetamido-2-deoxy-alpha-D-glucopyranose, with Beta 1-4 linkages connecting the four monosaccharide units. We use PDB ID 1LMN [29] as the unbound conformation of the protein, while the conformation of the glycan was generated with the GLYCAM webserver (https://glycam.org [30]). The protein closely resembles its bound form, with a backbone RMSD of 0.46 Å, while the modeled oligosaccharide does not fully align with the solved bound structure, especially in the ring conformation of 2-acetamido-2-deoxy-alpha-D-glucopyranose with a heavy atom RMSD between the two glycan conformations of 1.35 Å. Ambiguous interaction restraints are defined between the interface amino acids of the protein (defined as active) and the entire glycan (defined as passive).

**Figure 3:**
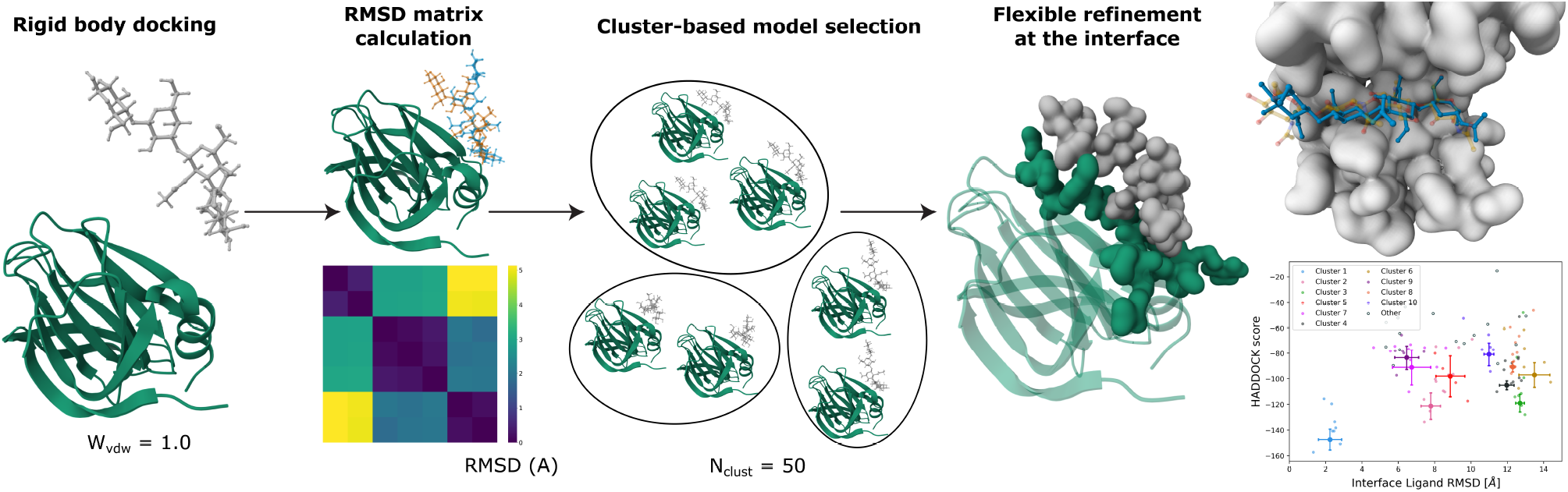
Workflow scheme for the protein-glycan modelling pipeline with a clustering step after rigid-body docking. The models that are chosen in this cluster-based selection are then subjected to a flexible refinement of the interface, and again clustered and analyzed. Figure adapted from Ref. [27].

When running the standard HADDOCK workflow, the best-scored docking models display values of DockQ around 0.47, with Interface-ligand RMSD (IL-RMSD) between 6 and 7 Å. The new docking workflow where RMSD clustering is performed between rigid-body docking and flexible refinement stages gives significantly better results, with the best cluster showing a mean DockQ of 0.83 and a mean IL-RMSD equal to 2.24 Å in its top four models. These markedly improved results are due to the clustering step, which performs a selection based on conformational diversity rather than purely selecting models according to the HADDOCK scoring function. Indeed, at the rigid-body docking step, the accurate models are not ranked among the top 200 in this case, with the best model (DockQ = 0.70) being ranked in position 947 out of 1000 sampled structures.

This example highlights that adding intermediate clustering steps and performing a cluster-based selection of docking models can significantly improve the docking results at the end of the workflow, especially when one or more input molecules are not modeled with high accuracy.

### Consensus scoring

Scoring models of biomolecular complexes is one of the core tasks of an integrative modelling platform, as it allows, if successful, to discriminate between near-native and non-native complexes. The scoring experiment has been included in the CAPRI challenge since 2005 [31]. In this context, research groups are asked to rank an ensemble of typically a few thousand models for each complex, to select the best 5 or 10 models. The scoring is considered successful if at least one good model, according to CAPRI criteria, is present in this reduced set.

A plethora of scoring functions have been developed over the recent years [32, 33, 34, 35, 36], each one with its own strengths and weaknesses.

Applying multiple scoring functions to the same set of complexes can be beneficial to improve ranking accuracy [34, 37, 38]. The HADDOCK3 suite is perfect for this task, as its modularity allows users to fine-tune scoring workflows by sequentially applying various types of scoring functions and inspecting the final consensus results. Five scoring modules are already available in the software, with the possibility to easily add more of them by exploiting HADDOCK3’s modularity. Additional clustering and analysis steps may be added to such workflows to group similar models together and to visualize their features.

We tested this approach in CAPRI round 57, where the HADDOCK3 energy minimisation and scoring were successfully combined with the VoroIF-jury method [34]. Here we focus on Target 268, an antibody-peptide complex, for which HADDOCK3 not only submitted the best model across the whole CAPRI scoring set (DockQ = 0.72, high-quality model according to CAPRI criteria) but also ranked it in the first position of its scoring ensemble. Other two scorer groups (Olechnovic [34] and LZERD [39]) were able to submit a high-quality model in the reduced scoring set of five complexes, although with a slightly lower quality (DockQ = 0.69).

Fig. 4a shows the consensus scoring procedure performed throughout CAPRI round 57. The models to be scored are provided as a single PDB file containing an ensemble of models (with the MODEL/ENDMDL construction). The workflow begins with building the topology of the input models (`topoaa`), followed by a short energy minimization step after which the HADDOCK scoring function is applied (`emscoring`). Next, FCC clustering [21] (`clustfcc` module) is performed and the top four models of the best ten or fifteen clusters are selected for the final evaluation (`seletopclusts`). Finally, the VoroIF-jury method (embedded in the `voroscoring` module) is applied to provide an alternative scoring function. In Fig. 4b we show the relationship between the two scoring functions applied to CAPRI target 268: according to VoroIF-jury, both cluster 1 and cluster 4 are reasonable choices, but the HADDOCK scoring function helps remove this ambiguity by assigning a significantly better score to cluster 1.

**Figure 4:**
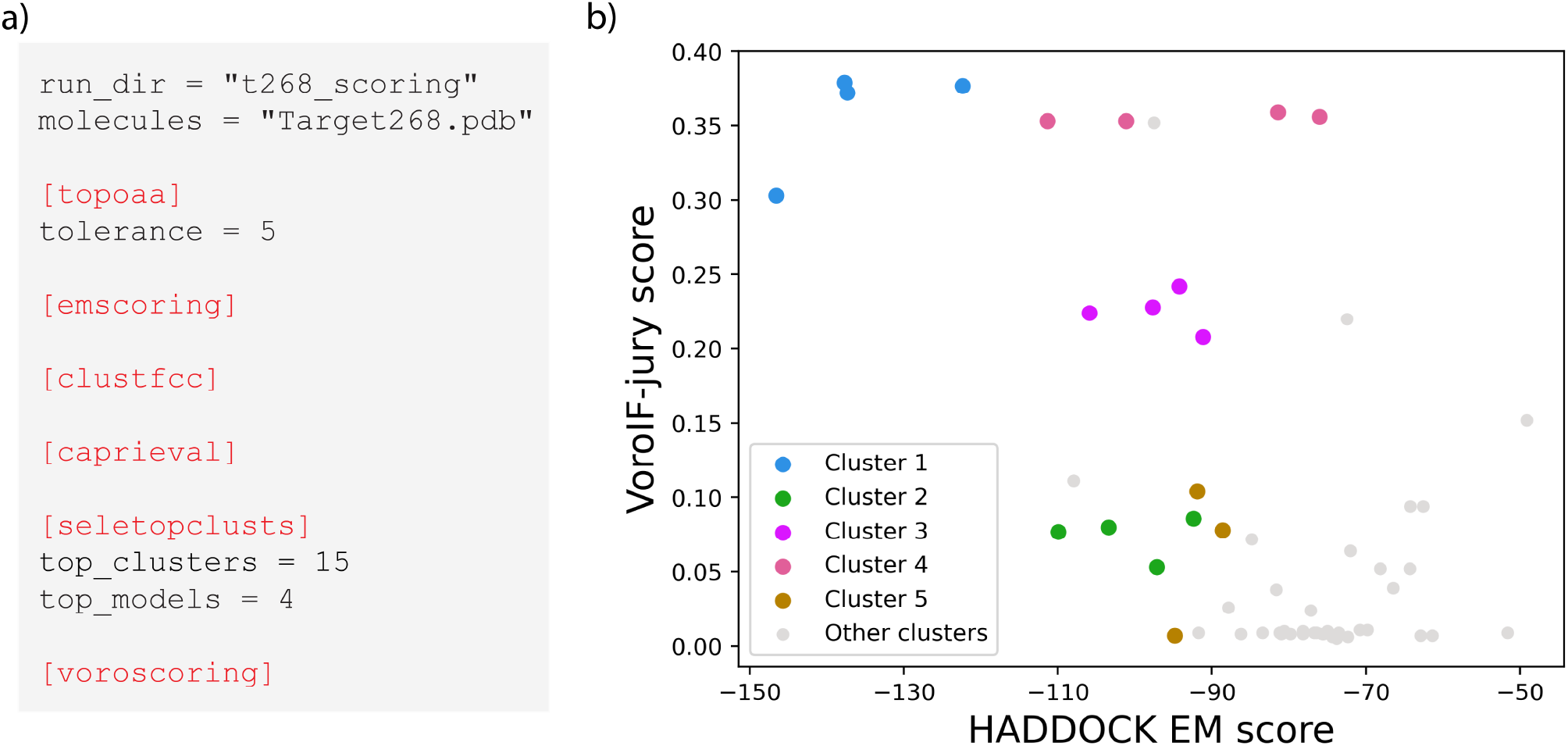
Illustration of HADDOCK3 scoring. a) The HADDOCK3 scoring workflow executed during CAPRI round 57. b) scatter plot between HADDOCK EM score (x axis) and VoroIF-jury score (y axis) for CAPRI target 268. The four models belonging to the top five FCC clusters are highlighted in colour, while models that are part of other clusters (from cluster 6 to cluster 15) are shown in gray. Models from cluster 1 stand out unambiguously in the top left corner (best - most negative - HADDOCK score and highest VoroIF-jury score). The best model in terms of energetics has a slightly lower VoroIF-jury score than the other cluster members.

This is only one example of the possibility to develop advanced consensus scoring workflows within the modular HADDOCK3 framework, for example combining general methods, such as the energy-based HADDOCK score, with functions more tailored to specific biomolecular complexes.

### Analysis of complexes and detection of hot spots

The analysis of a complex, either experimentally determined or predicted, can bring useful information about the residues involved in the interaction, to detect hot spots, and predict the potential impact of point mutations. In HADDOCK3, several analysis modules are available to inspect and describe biomolecular interactions. Among them, the contact-map (`contactmap`) module is dedicated to analysing contacts present in complexes and rendering them as both heat maps and chord charts (Fig. 5c and d). Interactive plots are generated using the *plotly* library [40], enabling a smooth navigation and understanding of the contacts in a given complex. In the case of a HADDOCK workflow with a clustering step, one plot per cluster will be generated, allowing for a fast identification of the different contact patches belonging to the various clusters. Another useful feature is provided by the alanine scanning (`alascan`) module, which detects residues at the interface of a complex and iteratively mutates them to alanine (default, but other mutations can be specified). The `alascan` module then performs a short energy minimisation and computes the HADDOCK score of the mutated complexes. For each amino acid at the interface (defined using a 5Å cutoff between any heavy atom of residues belonging to different chains), the module reports the difference in HADDOCK score between the wild-type sequence and the mutant, as well as the contribution of each component of the score (van der Waals, electrostatics, and desolvation energies). Once again, an interactive plot can be generated to obtain a graphical representation of the various energetical contributions of the interface residues (Fig. 5b) and possibly identify hot-spots. Although this module holds the name `alascan`, the conversion to any other amino acid is possible, thus providing a physics-based method to screen mutations in protein design projects.

**Figure 5:**
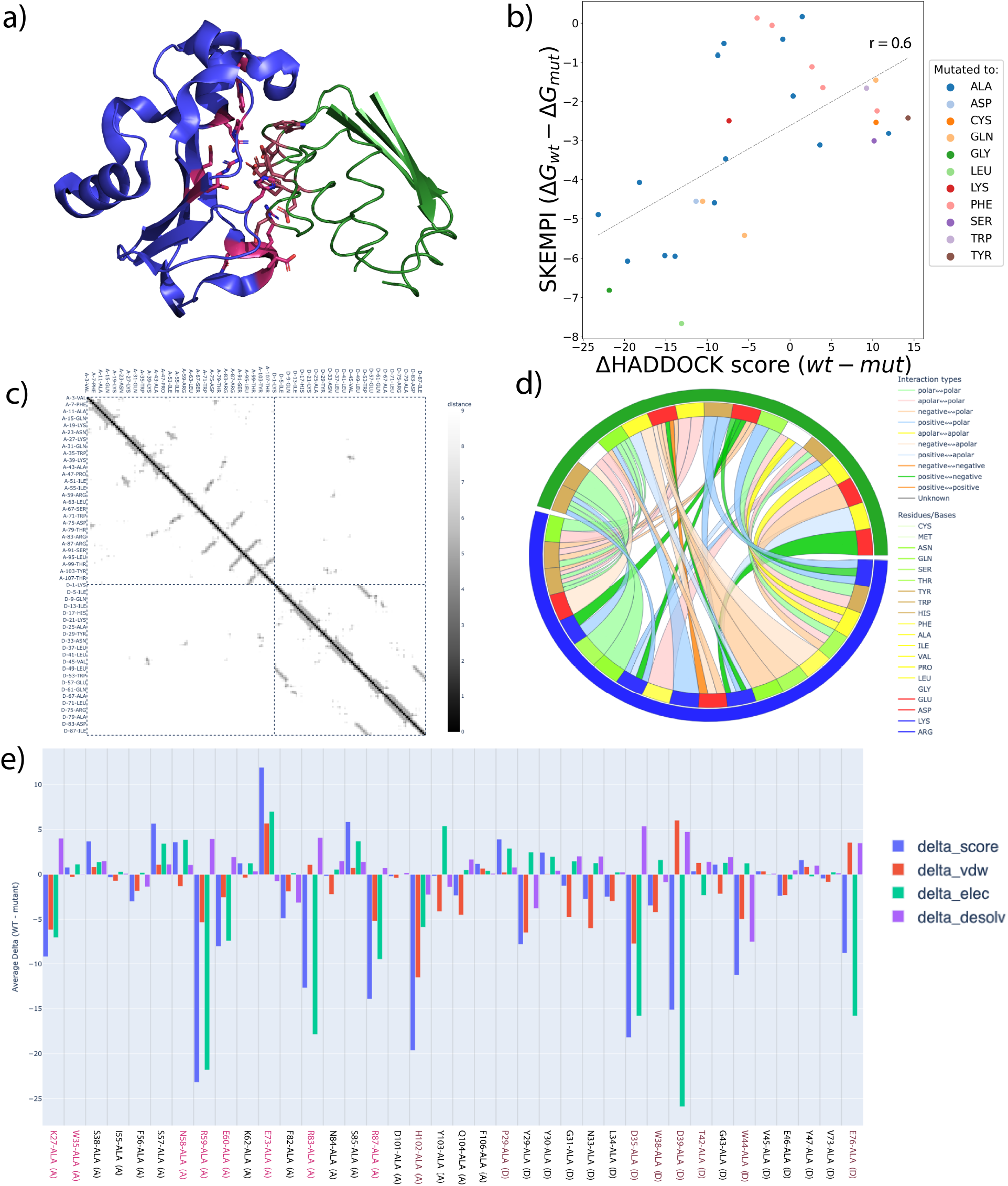
Interaction analysis with HADDOCK3. a) PyMOL [41] representation of the barnase-barstar complex (PDB ID 1BRS), where residues mutated present in the SKEMPI database are colored in red. b) Scatter plot of the difference in HADDOCK score (score wild-type - score mutant) obtained using the `alascan` module and difference in Gibbs free energy from SKEMPI (Δ*G*_*wt*_ − Δ*G*_*mut*_) on the barnase-barstar complex for single point mutations only. A Pearson correlation coefficient of 0.6 is obtained for this complex. c) Heat map, generated by the `contactmap` module, of the contacts observed on the PDB structure 1BRS. d) Chord chart, generated by the `contactmap` module of the inter-chains contacts observed on the PDB structure 1BRS. e) `alascan` plot displaying the influence of interface residues mutated to alanine. Colored labels are present in the SKEMPI dataset.

Here we show a combination of these analysis tools applied to the barnase-barstar complex (PDB ID 1BRS [42]) using the workflow shown in Fig. 6.

**Figure 6:**
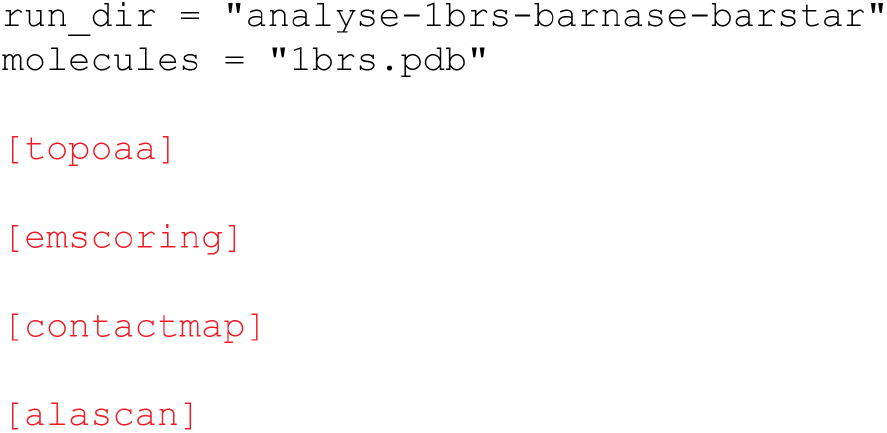
Example workflow for the analysis of a complex. Here the input complex (1brs.pdb) is provided as a single molecule and subjected to a short energy minimisation. The `contactmap` and `alascan` modules perform contact-map and alanine scanning analysis of the model, respectively (see main text).

After a short minimisation of the crystal structure, the intra and inter chains contacts are visualized as a heat map (Fig. 5c) by running the `contactmap` module, which also generates a chord chart coloring the contacts by type (Fig. 5d). Finally, the `alascan` module performs a systematic mutagenesis to Alanine of all interface residues. The resulting changes in HADDOCK score and its components are plotted (Fig. 5e). For this system, experimental binding affinity data are available from the SKEMPI database [43]. We extracted all single point values (ddG) and compared those to the changes in HADDOCK score from the `alascan` module (Fig. 5b). In this case, and without any optimisation, we observe a Pearson correlation of 0.6 between the experimental ddG and difference in HADDOCK score, showing that the HADDOCK score, together with the `alascan` module, can be used to obtain information about the influence of a given residue mutation. Note however that these observations are rather specific to the barnase-barstar complex, which has quite a hydrophilic/polar interface. From experience, correlations are expected to be worse for more hydrophobic interfaces. We refer to our previous work on predicting changes in binding affinities upon mutations for a more systematic analysis [44, 45].

## Conclusions

In this work we have introduced HADDOCK3, the new version of the integrative modelling software HADDOCK, which has been redesigned and modularized to provide higher flexibility and versatility in creating custom, system-specific modelling workflows. HADDOCK3 can seamlessly complement and assist machine learning-based structure prediction methods by incorporating available experimental data and physico-chemical features of the studied systems. A few possible applications of the method have been showcased, such as the flexible modelling of antibody-antigen complexes targeting multiple potential binding sites, clustering-enhanced protein-glycan interaction prediction, the consensus scoring of user-provided structural ensembles, and the analysis of intermolecular interactions.

HADDOCK3 comes with several external online resources facilitating the training of new users with tutorials (www.bonvinlab.org/education/HADDOCK3), tracking of potential coderelated issues (https://github.com/haddocking/haddock3/issues), providing day-to-day support to users (`ask.bioexcel.eu`) and accessing both the user manual (www.bonvinlab.org/haddock3-user-manual) and code documentation (www.bonvinlab.org/haddock3). In addition, on a yearly basis, a survey is conducted, where users are asked to give feedback and can request new features. This provides a basis for defining guidelines for the implementation of future features.

In conclusion, HADDOCK3, now easily available as a python package through PyPI, is a versatile and comprehensive software suite offering a wide range of options to address complex challenges in structural biology. We foresee that its new modular architecture will attract external software developers and users to contribute new modules and share workflows.

## Acknowledgements

The authors acknowledge the HADDOCK and BioExcel communities (https://bioexcel.eu) for providing constructive feedback during the process of software development and testing.

## Author contributions

M.G., V.R. and A.B. conceived the use cases and wrote the manuscript. M.G. and V.R. planned and ran the use cases and analysed the data. M.G., V.R., J.T., B.J-G, R.H., A.B. contributed to the design and implementation of the source code. M.G., V.R., R.H., A.K., X.X., R.V., A.E., S.V., and A.B. provided suggestions and feedback on the source code and contributed to the tutorials and documentation. A.B. supervised the project.

## Data Availability

Data, scripts, and HADDOCK3 configuration files used and described in this paper are available online on a dedicated GitHub repository hosted at https://github.com/haddocking/haddock3-paper-data

## References

[1] J. Jumper, R. Evans, A. Pritzel, T. Green, M. Figurnov, O. Ronneberger, K. Tunyasuvunakool, R. Bates, A. Žídek, A. Potapenko, et al., “Highly accurate protein structure prediction with alphafold,” nature, vol. 596, no. 7873, pp. 583–589, 2021.

[2] R. Evans, M. O’Neill, A. Pritzel, N. Antropova, A. Senior, T. Green, A. Žídek, R. Bates, S. Blackwell, J. Yim, et al., “Protein complex prediction with alphafold-multimer,” biorxiv, pp. 2021–10, 2021.

[3] M. Baek, F. DiMaio, I. Anishchenko, J. Dauparas, S. Ovchinnikov, G. R. Lee, J. Wang, Q. Cong, L. N. Kinch, R. D. Schaeffer, C. Millán, H. Park, C. Adams, C. R. Glassman, A. DeGiovanni, J. H. Pereira, A. V. Rodrigues, A. A. van Dijk, A. C. Ebrecht, D. J. Opperman, T. Sagmeister, C. Buhlheller, T. Pavkov-Keller, M. K. Rathinaswamy, U. Dalwadi, C. K. Yip, J. E. Burke, K. C. Garcia, N. V. Grishin, P. D. Adams, R. J. Read, and D. Baker, “Accurate prediction of protein structures and interactions using a three-track neural network,” Science, vol. 373, no. 6557, pp. 871–876, 2021.

[4] R. Wu, F. Ding, R. Wang, R. Shen, X. Zhang, S. Luo, C. Su, Z. Wu, Q. Xie, B. Berger, et al., “High-resolution de novo structure prediction from primary sequence,” BioRxiv, pp. 2022–07, 2022.

[5] Z. Lin, H. Akin, R. Rao, B. Hie, Z. Zhu, W. Lu, N. Smetanin, R. Verkuil, O. Kabeli, Y. Shmueli, A. dos Santos Costa, M. Fazel-Zarandi, T. Sercu, S. Candido, and A. Rives, “Evolutionary-scale prediction of atomic-level protein structure with a language model,” Science, vol. 379, no. 6637, pp. 1123–1130, 2023.

[6] M. Baek, R. McHugh, I. Anishchenko, H. Jiang, D. Baker, and F. DiMaio, “Accurate prediction of protein–nucleic acid complexes using rosettafoldna,” Nature methods, vol. 21, no. 1, pp. 117–121, 2024.

[7] R. Krishna, J. Wang, W. Ahern, P. Sturmfels, P. Venkatesh, I. Kalvet, G. R. Lee, F. S. Morey-Burrows, I. Anishchenko, I. R. Humphreys, et al., “Generalized biomolecular modeling and design with rosettafold all-atom,” Science, vol. 384, no. 6693, p. eadl2528, 2024.

[8] J. Abramson, J. Adler, J. Dunger, R. Evans, T. Green, A. Pritzel, O. Ronneberger, L. Willmore, A. J. Ballard, J. Bambrick, et al., “Accurate structure prediction of biomolecular interactions with alphafold 3,” Nature, pp. 1–3, 2024.

[9] R. V. Honorato, M. E. Trellet, B. Jiménez-García, J. J. Schaarschmidt, M. Giulini, V. Reys, P. I. Koukos, J. P. Rodrigues, E. Karaca, G. C. van Zundert, et al., “The haddock2. 4 web server for integrative modeling of biomolecular complexes,” Nature protocols, vol. 19, no. 11, pp. 3219–3241, 2024.

[10] V. Reys, M. Giulini, V. Cojocaru, A. Engel, X. Xu, J. Roel-Touris, C. Geng, F. Ambrosetti, B. Jiménez-García, Z. Jandova, et al., “Integrative modeling in the age of machine learning: A summary of haddock strategies in capri rounds 47–55,” Proteins: Structure, Function, and Bioinformatics, 2024.

[11] C. Dominguez, R. Boelens, and A. M. Bonvin, “Haddock: a protein-protein docking approach based on biochemical or biophysical information,” Journal of the American Chemical Society, vol. 125, no. 7, pp. 1731–1737, 2003.

[12] B. Jiménez-García, J. Roel-Touris, M. Romero-Durana, M. Vidal, D. Jiménez-González, and J. Fernández-Recio, “Lightdock: a new multi-scale approach to protein–protein docking,” Bioinformatics, vol. 34, no. 1, pp. 49–55, 2018.

[13] S. Feng, Z. Chen, C. Zhang, Y. Xie, S. Ovchinnikov, Y. Q. Gao, and S. Liu, “Integrated structure prediction of protein–protein docking with experimental restraints using colabdock,” Nature Machine Intelligence, vol. 6, no. 8, pp. 924–935, 2024.

[14] S. J. De Vries, A. D. Van Dijk, M. Krzeminski, M. van Dijk, A. Thureau, V. Hsu, T. Wassenaar, and A. M. Bonvin, “Haddock versus haddock: new features and performance of haddock2. 0 on the capri targets,” Proteins: structure, function, and bioinformatics, vol. 69, no. 4, pp. 726–733, 2007.

[15] G. Van Zundert, J. Rodrigues, M. Trellet, C. Schmitz, P. Kastritis, E. Karaca, A. Melquiond, M. van Dijk, S. De Vries, and A. Bonvin, “The haddock2. 2 web server: user-friendly integrative modeling of biomolecular complexes,” Journal of molecular biology, vol. 428, no. 4, pp. 720–725, 2016.

[16] P. Eastman and V. Pande, “Openmm: A hardware-independent framework for molecular simulations,” Computing in science & engineering, vol. 12, no. 4, pp. 34–39, 2010.

[17] A. Vangone, J. Schaarschmidt, P. Koukos, C. Geng, N. Citro, M. E. Trellet, L. C. Xue, and A. M. Bonvin, “Large-scale prediction of binding affinity in protein–small ligand complexes: the prodigy-lig web server,” Bioinformatics, vol. 35, no. 9, pp. 1585–1587, 2019.

[18] P. L. Kastritis, J. P. Rodrigues, G. E. Folkers, R. Boelens, and A. M. Bonvin, “Proteins feel more than they see: fine-tuning of binding affinity by properties of the non-interacting surface,” Journal of molecular biology, vol. 426, no. 14, pp. 2632–2652, 2014.

[19] A. Vangone and A. M. Bonvin, “Contacts-based prediction of binding affinity in protein– protein complexes,” elife, vol. 4, p. e07454, 2015.

[20] J. Janin, K. Henrick, J. Moult, L. T. Eyck, M. J. Sternberg, S. Vajda, I. Vakser, and S. J. Wodak, “Capri: a critical assessment of predicted interactions,” Proteins: Structure, Function, and Bioinformatics, vol. 52, no. 1, pp. 2–9, 2003.

[21] J. P. Rodrigues, M. Trellet, C. Schmitz, P. Kastritis, E. Karaca, A. S. Melquiond, and A. M. Bonvin, “Clustering biomolecular complexes by residue contacts similarity,” Proteins: Structure, Function, and Bioinformatics, vol. 80, no. 7, pp. 1810–1817, 2012.

[22] S. Basu and B. Wallner, “Dockq: a quality measure for protein-protein docking models,” PloS one, vol. 11, no. 8, p. e0161879, 2016.

[23] M. Giulini, R. V. Honorato, J. L. Rivera, and A. M. Bonvin, “Arctic-3d: automatic retrieval and clustering of interfaces in complexes from 3d structural information,” Communications Biology, vol. 7, no. 1, p. 49, 2024.

[24] M. Blech, D. Peter, P. Fischer, M. M. Bauer, M. Hafner, M. Zeeb, and H. Nar, “One target—two different binding modes: structural insights into gevokizumab and canakinumab interactions to interleukin-1β,” Journal of molecular biology, vol. 425, no. 1, pp. 94–111, 2013.

[25] M. Varadi, J. Berrisford, M. Deshpande, S. S. Nair, A. Gutmanas, D. Armstrong, L. Pravda, B. Al-Lazikani, S. Anyango, G. J. Barton, et al., “Pdbe-kb: a community-driven resource for structural and functional annotations,” Nucleic Acids Research, vol. 48, no. D1, pp. D344–D353, 2020.

[26] A. Varki, R. D. Cummings, J. D. Esko, P. Stanley, G. W. Hart, M. Aebi, A. G. Darvill, T. Kinoshita, N. H. Packer, J. H. Prestegard, et al., “Essentials of glycobiology [internet],” 2015.

[27] A. Ranaudo, M. Giulini, A. Pelissou Ayuso, and A. M. Bonvin, “Modeling protein–glycan interactions with haddock,” Journal of Chemical Information and Modeling, vol. 64, no. 19, pp. 7816–7825, 2024.

[28] S. Karlsen and E. Hough, “Crystal structures of three complexes between chito-oligosaccharides and lysozyme from the rainbow trout. how distorted is the nag sugar in site d?,” Biological Crystallography, vol. 51, no. 6, pp. 962–978, 1995.

[29] S. Karlsen, B. E. Eliassen, L. K. Hansen, R. L. Larsen, B. W. Riise, A. Smalås, E. Hough, and B. Grinde, “Refined crystal structure of lysozyme from the rainbow trout (oncorhynchus mykiss),” Biological Crystallography, vol. 51, no. 3, pp. 354–367, 1995.

[30] “Glycam web-server.” http://glycam.org/, 2015.

[31] M. F. Lensink, R. Méndez, and S. J. Wodak, “Docking and scoring protein complexes: Capri 3rd edition,” Proteins: Structure, Function, and Bioinformatics, vol. 69, no. 4, pp. 704–718, 2007.

[32] N. Renaud, C. Geng, S. Georgievska, F. Ambrosetti, L. Ridder, D. F. Marzella, M. F. Réau, A. M. Bonvin, and L. C. Xue, “Deeprank: a deep learning framework for data mining 3d protein-protein interfaces,” Nature communications, vol. 12, no. 1, p. 7068, 2021.

[33] J. P. Roney and S. Ovchinnikov, “State-of-the-art estimation of protein model accuracy using alphafold,” Physical Review Letters, vol. 129, no. 23, p. 238101, 2022.

[34] K. Olechnovič, L. Valančauskas, J. Dapkūnas, and Č. Venclovas, “Prediction of protein assemblies by structure sampling followed by interface-focused scoring,” Proteins: Structure, Function, and Bioinformatics, vol. 91, no. 12, pp. 1724–1733, 2023.

[35] P. Bryant, G. Pozzati, and A. Elofsson, “Improved prediction of protein-protein interactions using alphafold2,” Nature communications, vol. 13, no. 1, p. 1265, 2022.

[36] M. Réau, N. Renaud, L. C. Xue, and A. M. Bonvin, “Deeprank-gnn: a graph neural network framework to learn patterns in protein–protein interfaces,” Bioinformatics, vol. 39, no. 1, p. btac759, 2023.

[37] S. Liang, S. O. Meroueh, G. Wang, C. Qiu, and Y. Zhou, “Consensus scoring for enriching near-native structures from protein–protein docking decoys,” Proteins: Structure, Function, and Bioinformatics, vol. 75, no. 2, pp. 397–403, 2009.

[38] N. Manshour, J. Z. Ren, F. Esmaili, E. Bergstrom, and D. Xu, “Comprehensive evaluation of alphafold-multimer, alphafold3 and colabfold, and scoring functions in predicting protein-peptide complex structures,” bioRxiv, pp. 2024–11, 2024.

[39] V. Venkatraman, Y. D. Yang, L. Sael, and D. Kihara, “Protein-protein docking using region-based 3d zernike descriptors,” BMC bioinformatics, vol. 10, pp. 1–21, 2009.

[40] “Plotly technologies inc., collaborative data science,” 2015.

[41] Schrödinger, LLC, “The PyMOL molecular graphics system, version 1.8.” November 2015.

[42] A. M. Buckle, G. Schreiber, and A. R. Fersht, “Protein-protein recognition: Crystal structural analysis of a barnase-barstar complex at 2.0-. ang. resolution,” Biochemistry, vol. 33, no. 30, pp. 8878–8889, 1994.

[43] J. Jankauskaite, B. Jiménez-García, J. Dapkūnas, J. Fernández-Recio, and I. H. Moal, “Skempi 2.0: an updated benchmark of changes in protein–protein binding energy, kinetics and thermodynamics upon mutation,” Bioinformatics, vol. 35, no. 3, pp. 462–469, 2019.

[44] C. Geng, L. C. Xue, J. Roel-Touris, and A. M. Bonvin, “Finding the δδg spot: Are predictors of binding affinity changes upon mutations in protein–protein interactions ready for it?,” Wiley Interdisciplinary Reviews: Computational Molecular Science, vol. 9, no. 5, p. e1410, 2019.

[45] C. Geng, Y. Jung, N. Renaud, V. Honavar, A. M. Bonvin, and L. C. Xue, “iscore: a novel graph kernel-based function for scoring protein–protein docking models,” Bioinformatics, vol. 36, no. 1, pp. 112–121, 2020.

